# Corruption of DNA End-Joining in Mammalian Chromosomes by Progerin Expression as Revealed by a Model Cell Culture System

**DOI:** 10.1101/2022.07.18.500488

**Authors:** Liza A. Joudeh, Alannah J. DiCintio, Madeline R. Ries, Andrew S. Gasperson, Kennedy E. Griffin, Victoria P. Robbins, Makenzie Bonner, Sarah Nolan, Emma Black, Alan S. Waldman

## Abstract

Hutchinson-Gilford Progeria Syndrome (HGPS) is a rare genetic condition characterized by features of accelerated aging and a life expectancy of about 14 years. HGPS is commonly caused by a point mutation in the LMNA gene which codes for lamin A, an essential component of the nuclear lamina. The HGPS mutation alters splicing of the LMNA transcript, leading to a truncated, farnesylated form of lamin A termed “progerin.” HGPS is associated with accumulation of genomic DNA double-strand breaks (DSBs), suggesting altered DNA repair. DSB repair normally occurs by either homologous recombination (HR), an accurate, templated form of repair, or by non-homologous end joining (NHEJ), an error-prone non-templated rejoining of DNA ends. Some NHEJ events occur via high-fidelity joining of DNA ends and we refer to such events as precise ligation (PL). Previously, we reported that expression of progerin correlated with increased NHEJ relative to HR. We now report on progerin’s impact on the nature of DNA end-joining. We used a model system involving a DNA end-joining reporter substrate integrated into the genome of cultured thymidine kinase-deficient mouse fibroblasts. Some cells were engineered to express progerin. DSBs were induced in the substrate through expression of endonuclease I-SceI, and DSB repair events were recovered through selection for thymidine kinase function. Progerin expression correlated with a significant shift away from PL and toward error-prone NHEJ. Our work suggests that progerin suppresses interactions between complementary sequences at DNA termini, shifting DSB repair toward low-fidelity DNA end-joining and perhaps contributing to aging through compromised genome stability.

## INTRODUCTION

In the face of daily damage to DNA, the ability of mammalian cells to maintain a stable genome is dependent on the functionality of a variety of DNA repair pathways. Among the various forms of damage that must be dealt with is the DNA double-strand break (DSB). DSBs can be generated spontaneously in numerous ways, including processing of a variety of DNA lesions and at stalled or collapsed replication forks. DSBs can also be induced by exposure to certain chemicals or radiation. Regardless of their origin, it is essential that DSBs be repaired quickly and reasonably accurately to avoid potentially deleterious chromosomal rearrangements or mutations. Genetic variations that impinge on DSB repair processes would thus be expected to impact cellular and organismal viability. Conversely, a genetic disorder that engenders reduced viability may be associated with faulty DNA repair. In our current work, we used a model cell culture system to explore how the rare laminopathy known as Hutchinson-Gilford Progeria Syndrome (HGPS) [reviewed in 1], a premature aging disorder, may alter the nature of DSB repair.

To mend DSBs, mammalian cells utilize two types of repair pathways: homologous recombination (HR) and non-homologous end-joining (NHEJ) [reviewed in 2-11]. Although there are numerous sub-pathways for HR and NHEJ, the critical difference between these repair strategies is that HR utilizes a template sequence to maintain or restore genetic information at the DSB site that may otherwise be altered, whereas NHEJ involves no template in the rejoining of DNA ends. HR is thus generally considered to be accurate while NHEJ often produces deletions, or sometimes insertions, and, thus, is generally error-prone. Additionally, HR is active primarily during the late S or G2 stage of the cell cycle in dividing cells, whereas NHEJ is active throughout the cell cycle and in non-dividing cells.

Lack of change is the essence of stability. For a mammalian genome to truly remain stable, DNA repair pathways must be executed with high accuracy. With regards to DSB repair it would seem, therefore, that HR underpins genome stability since HR intrinsically provides greater accuracy than NHEJ. However, the potential exists for HR-generated genomic instability should HR occur at abnormally high levels, which may lead to increased loss-of-heterozygosity of deleterious alleles [12,13]. Genetic perturbations that allow the choice of inappropriate recombination partners can also lead to genomic instability in the form of localized sequence alterations or gross chromosomal rearrangements [11,14,15]. Crossovers allowed to occur between imperfectly matched sequences may lead to translocations. To avoid mutation or rearrangements, it is important that genetic exchange only occurs among sequences that share perfect or near-perfect homology. Along these lines, mammalian cells normally exert stringent control over HR, allowing exchange to occur only between those sequences that exhibit a very high degree of sequence identity [16-18].

As noted, NHEJ is generally considered to be error-prone, often producing sequence deletions or insertions. Minimization of the size of NHEJ-associated deletions or insertions would likely serve to lessen the impact of NHEJ on cellular functions and, indeed, a portion of end-joining events proceed with no alteration at all to the sequences at the break site [19]. We specifically refer to this latter class of events as precise ligation (PL). The size of NHEJ-associated sequence deletions or insertions, the balance between error-prone NHEJ versus PL, along with the appropriate regulation of HR are all germane to the maintenance of genome stability.

The consequences of the failure of DNA transactions to maintain genome stability are varied. Aberrant HR or DSB repair is often associated with cancer [20-22]. Genomic instability is also viewed as a possible basis for, or at least a significant contributor to, the aging process. Increased levels of DNA damage, mutation, and large-scale chromosomal alterations including translocations, insertions and deletions have been observed with increasing age in humans and other organisms [23-36]. As genome integrity is diminished over time, critical cellular functions would be expected to be disturbed. In addition, cell number would gradually be reduced as cells are lost due to apoptotic responses to unrepaired DNA lesions, leading to tissue depletion and loss of biological functions. Alternatively, disruption of stress responses, including DNA damage responses, can lead to uncontrolled cellular division in the form of cancerous growth rather than cellular death, and cancer is indeed often viewed as a disease of aging. The links between genomic instability in both aging and cancer, two conditions that superficially seem to be at odds on the cellular level, are thereby overlapping.

Much evidence has been reported that the increase in genomic instability that accompanies aging correlates with a decrease in the intrinsic efficiency of or alteration of the nature of a variety of DNA repair pathways as a function of age [23-36]. Alterations in NHEJ were reported in rat brain during aging [29], and studies with mice have suggested that the fidelity of DSB repair diminishes with age [30]. Both the efficiency and fidelity of DSB repair has been observed to decrease as human fibroblasts approach senescence [31]. Chromosomal DSBs accumulate in human cells approaching senescence, and it has been suggested that DSBs may be involved directly in the actual induction of senescence [23]. In short, evidence abounds for a role for impaired or altered repair of DSBs in the biology of aging.

Genetic disorders that produce clinical features of premature aging (progeria) are often associated with innate DNA repair defects and associated genomic instability [24-28, 32-36]. Hutchinson–Gilford Progeria Syndrome (HGPS) is one such rare genetic syndrome that leads to accelerated aging [1]. The average lifespan of an individual with HGPS is about fourteen years. The most common cause of HGPS is a point mutation in the LMNA gene which normally codes for lamin A and its splice variant lamin C. The mutation responsible for HGPS leads to increased usage of a cryptic splice site which leads to the production of a truncated form of lamin A referred to as “progerin.” Significantly, it has been learned that progerin is in fact expressed at low levels in healthy individuals and appears to play a role in the normal aging process [37-40]. Unlike wild-type fully processed lamin A, progerin retains a farnesyl group at its carboxy terminus. This farnesyl group causes progerin to largely remain associated with the inner nuclear membrane rather than localize to the nuclear lamina where lamin A normally resides.

Lamin A serves as an important component of the nuclear lamina which plays structural as well as catalytic roles in the nucleus. In HGPS, the impact of progerin over-expression on nuclear architecture is severe. The nuclei of HGPS cells are characteristically misshapen and display blebs and invaginations, and this altered nuclear structure conveys changes to numerous nuclear functions. Progerin expression in HGPS interferes with recruitment of replications factors to replication forks and this leads to replication fork stalling and collapse [36,40-44]. A body of literature also implicates lamin A and its variants in DNA repair [45-51].

One important consequence of progerin expression in HGPS cells is an accumulation of DSBs and marked sensitivity to DNA damaging agents [41-44,52,53]. The general conclusion that has been drawn is that DSB repair in HGPS is delayed or perhaps sometimes precluded, and this corruption of DNA repair is related to reduced lifespan of cells and individuals. It has also been reported that recruitment of DSB repair proteins, particularly those involved in HR repair (including Rad 50, Rad51, NBS1, and MRE11), to sites of chromosomal damage is delayed in HGPS [41,53]. This delay in recruitment of HR-associated proteins presents an explanation for observations suggesting that NHEJ may be enhanced while HR is concomitantly reduced by progerin expression [47,48,50,54]. Recently, we confirmed these latter studies by directly demonstrating that progerin expression shifts repair pathway choice at a defined genomic DSB away from the HR pathway and towards NHEJ [55].

As noted above, the characteristics of NHEJ as well as the balance between PL versus mutagenic NHEJ events are all factors expected to contribute to the maintenance of genome stability. It is thus of considerable interest to explore whether progerin expression not only shifts DSB repair toward DNA end-joining, but whether progerin additionally alters the nature of DNA end-joining. In the current work, we used a novel model experimental system in mouse fibroblasts and show that progerin expression alters DNA end-joining by bringing about a marked decrease in the frequency of PL events with a concomitant increase in error-prone NHEJ. Our data also suggest that progerin expression correlates with a possible increase in deletion size associated with NHEJ. Additional experiments show that progerin expression does not allow genetic exchange between imperfectly matched sequences and thus, while reducing the occurrence or HR, progerin does not decrease the fidelity of HR. Collectively, our work suggests that progerin expression compromises genome stability by corrupting interactions between complementary sequences at DNA termini during DNA repair. Such influences of progerin may impact genome stability and contribute to the aging process.

## MATERIALS AND METHODS

### General cell culture

All cell lines were derived from Ltk^-^ mouse fibroblasts and were grown in Dulbecco’s Modified Eagle Medium (DMEM/low glucose) supplemented with 10% heat-inactivated fetal bovine serum, minimal essential medium non-essential amino acids, and gentamicin (50 μg/ml). Cells were maintained in a 37°C incubator in a 5% CO_2_ atmosphere.

### DNA repair substrates

Substrates pTKSce2 and pHOME are based on the vector pJS-1 [56,57], which is a derivative of pSV2neo [58].

pTKSce2, described previously [19], contains a defective copy of the herpes simplex virus type one thymidine kinase (tk) gene on a 2.5-kb BamHI fragment inserted into the unique BamHI site of the vector. The tk gene was rendered nonfunctional by insertion of a 47 bp oligonucleotide after position 963 of the tk gene coding region (numbering as described by Wagner et al. [59]). The 47 bp oligonucleotide contains two 18 bp I-SceI recognition sites. pTKSce2 was designed as a reporter of both PL as well as error-prone NHEJ.

Substrate pHOME, described previously [60], contains a “recipient” herpes simplex virus type one tk gene on a 2.5-kb BamHI fragment inserted into the BamHI site of the vector. The recipient tk gene in pHOME was rendered nonfunctional by insertion of a 30-bp oligonucleotide after nucleotide position 1,215 in the tk gene. The 30-bp oligonucleotide sequence includes the 18 bp recognition site for endonuclease I-SceI. Also included in pHOME, at the unique HindIII site in the vector, is a “donor” sequence that shares about 80% nucleotide identity (20% divergence) with the recipient. The donor sequence on pHOME is a nonfunctional EcoRV-StuI fragment (nucleotides 848–1,668) of the herpes simplex type 2 tk gene, which is missing 30% of the tk coding region at the 5′ end. As described [60], several nucleotide changes were made on the donor sequence so that the donor encodes an amino acid sequence identical to that encoded by the corresponding region of the recipient tk gene despite displaying about 20% sequence divergence with the recipient. Substrate pHOME was designed as a reporter or HeR and NHEJ [60].

### Recombination and DSB repair reporter cell lines

Substrate pTKSce2 was linearized by digestion with ClaI and stably transfected into mouse Ltk^-^ cells via electroporation as previously described [61]. Transfected clones were recovered by selection in medium containing G418 (200 μg/ml). Cell lines containing a single integrated copy of pTKSce2 were identified by Southern blotting analysis. One such cell line designated line “13” was used in subsequent experiments.

The isolation of cell line HOME-1-24 from mouse Ltk^-^ cells was described previously [60]. Cell line HOME 1-24 contains a single stably integrated copy of pHOME.

To produce derivatives of cell lines 13 and HOME-1-24 that stably express GFP-progerin, 5 × 10^6^ cells of each cell line were electroporated with a mixture of 10 μg of plasmid pEGFP-D50 lamin A (Addgene plasmid #17653) that had been linearized with EagI and 1 μg of pBABE-puro (Addgene plasmid #1764) which contains a puromycin resistance gene. Stably transfected clones were selected in medium containing puromycin (5 μg/ml). After 14 days of selection, colonies that showed nuclear GFP fluorescence were propagated further and GFP-progerin expression was confirmed by Western blot.

### Recovery of DSB repair events

To induce DSBs in cells containing an integrated DNA substrate, cells were electroporated with the I-SceI expression plasmid pCMV-3xnls-I-SceI (“pSce”) [61] as previously described [19]. Briefly, 5 × 10^6^ cells were mixed with 20 μg of pSce in a volume of 800 μL of phosphate buffered saline at room temperature and electroporated in a Bio-Rad Gene Pulser (Bio-Rad, Hercules, CA, USA) set to 750 V and 25 μFd. Following electroporation, cells were plated into growth medium at different densities under no selection and allowed to recover for two days. Cells were then fed with hypoxanthine/aminopterin/thymidine (HAT) medium [62] to select for tk^+^ clones. It was empirically determined that for cell line 13 and its derivatives, plating 5 × 10^4^ cells per 75 cm^2^ flask yielded well-separated colonies that were easily counted. Colonies were typically picked after 10-12 days of growth in HAT medium and propagated for further analysis.

### PCR amplification and DNA sequence analysis

A fragment of the tk gene spanning the original location of the gene-disrupting oligonucleotide was amplified from 500 ng of genomic DNA isolated from tk-positive clones using primers AW100 (5′-TAATACGACTCACTATAGGGTTGCGCCCTCGCCGGCAGC-3′) and AW133 (5′-CAGGAAACAGCTATGACCCGGTGGGGTATCGACAGAGT-3′). AW100 is composed of nucleotides 600-618 of the coding sequence of the HSV-1 tk gene (numbering according to [59]), with a T7 forward universal primer appended to the 5′ end of the primer. AW133 is composed of nucleotides 1786-1767 of the HSV-1 tk gene with an M13 reverse universal primer appended to the 5′ end of the primer. PCR was accomplished using Ready-To-Go PCR beads (Cytiva) and a “touchdown” protocol as previously described [63]. PCR products were then sequenced using a T7 primer by Eton Bioscience, Inc. (Research Triangle Park, NC).

### Southern blotting analysis

Genomic DNA samples (8 μg each) were digested with appropriate restriction enzymes and resolved on 0.8 % agarose gels. DNA was transferred to nitrocellulose membranes and hybridized with ^32^P-labeled *tk* probes as previously described [56].

### Western blots

Blots were performed using SDS-PAGE with 6% stacking gels and 8% separating gels. Each lane contained 30 μg of total cellular protein, and Bio-Rad Precision Plus Kaleidoscope Protein Standard (#161-0375, 5 μL) was used for molecular weight markers. Proteins were transferred to a nitrocellulose membrane. Ponceau S staining was performed for transfer efficiency and to confirm equal protein loading across lanes, and membranes were blocked in 2% non-fat milk in PBST (PBS plus 0.1% Tween 20) for two hours. The primary antibody used was GFP (B-2): sc-9996 (mouse monoclonal, from Santa Cruz Biotechnology, Inc.) at a dilution of 1:500 in 2% non-fat milk-PBST and incubated at 4 degrees Celsius overnight on a rocker. Primary antibody was removed after washing 3x with PBST for 10 minutes. Secondary antibody used was goat anti-mouse IgG-HRP: sc-2005 (Santa Cruz Biotechnology, Inc.) at a dilution of 1:1000 in 2% non-fat milk-PBST and incubated at room temperature for two hours.

The membrane was washed 3x with PBST prior to being developed. Detection was accomplished using GE Healthcare Amersham ECL Select Western Blotting Detection Reagent.

## RESULTS

### An experimental system for studying the impact of progerin expression on DNA end-joining in mammalian cells

To study NHEJ and PL within the genome of mammalian cells, mouse Ltk-cells were stably transfected with DNA substrate pTKSce2 (Figure 1) and a cell line containing a single integrated copy of pTKSce2 was isolated. This cell line was designated line “13.” As illustrated in Figure 1, pTKSce2 contains a tk gene that is disrupted by a 47 bp oligonucleotide that includes two recognition sites for endonuclease I-SceI. Endonuclease I-SceI makes a staggered DSB, with 4 bp single-stranded overhangs. If cells containing pTKSce2 are transfected with pSce (an I-SceI expression plasmid [61]), two DSBs can be induced. As shown in Figure 1, if DSBs are induced and the two outermost single-stranded overhangs are annealed and DNA ends joined with no other sequence alterations, then the 23 bp DNA segment between the two I-SceI breaks will be lost and a functional tk gene retaining the indicated 24 bp insert (which contains an I-SceI recognition site) will be produced. This clean ligation of DNA ends serves as a model PL event. The PL product will convert a cell from a tk^-^ phenotype to a tk^+^ phenotype and, thus, PL events can be recovered by selecting for HAT^R^ clones. Error-prone NHEJ events at one or both DSBs may also produce functional in-frame tk genes and these events can also be recovered under HAT selection following DSB induction.

**Figure 1.**
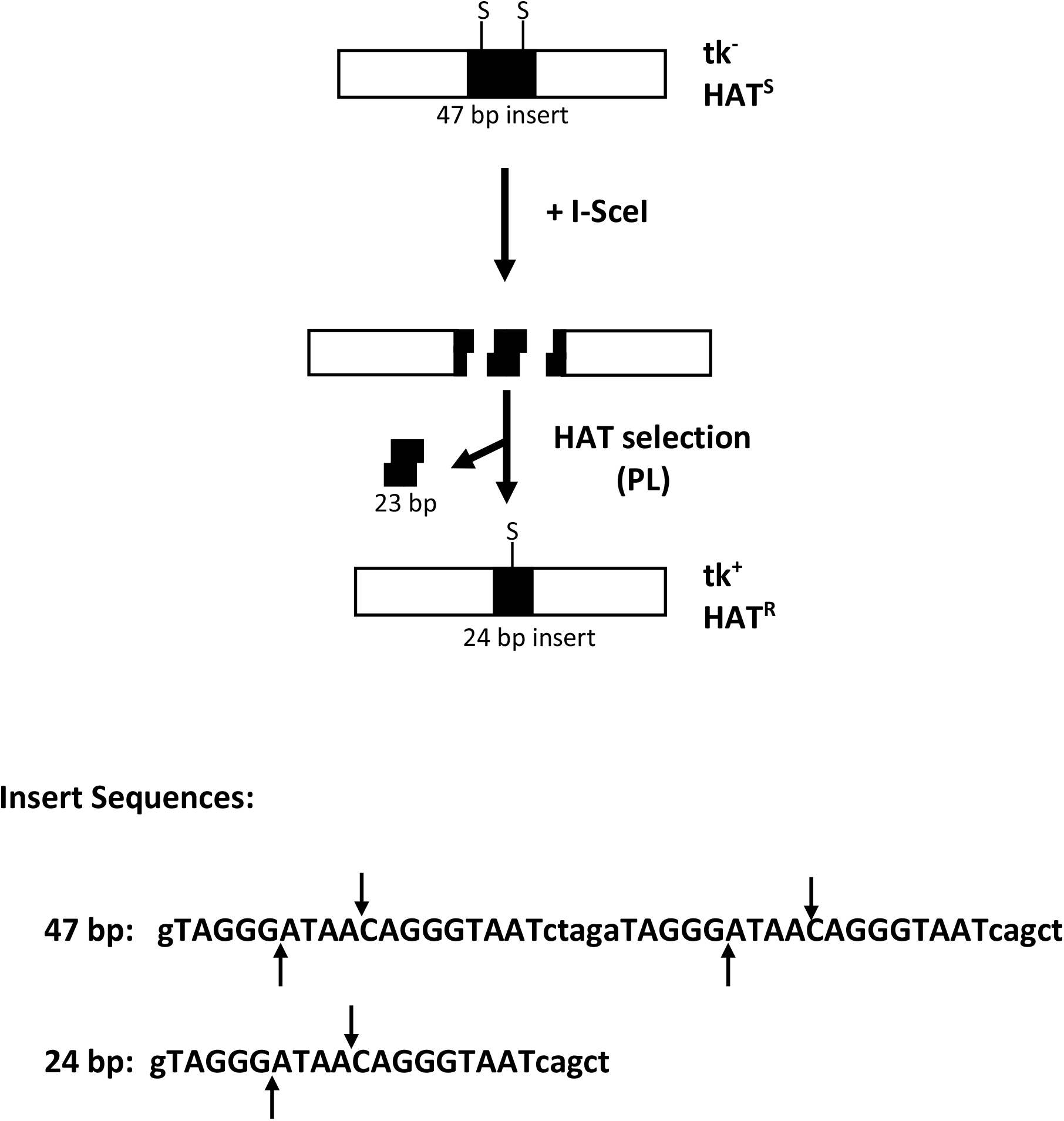
Strategy for recovering precise ligation (PL) events using DNA substrate pTKSce2. As represented schematically at the top of the figure, pTKSce2 contains a tk gene (white rectangle) disrupted by insertion of a 47 bp oligonucleotide containing two 18 bp I-SceI recognition sites (S). If endonuclease I-SceI is transiently expressed in a cell line containing an integrated copy of pTKSce2, two DSBs can be induced. A PL event involving joining of the two outermost sticky ends produced by I-SceI will produce a functional tk gene retaining a 24 bp insert containing an I-SceI site, and such an event is recoverable by HAT selection. The nucleotide sequences of the original 47 bp insert and the 24 bp insert produced by PL are illustrated. The 18 bp I-SceI recognition sequence is presented in uppercase font, and the actual sites of staggered cleavage by I-SceI are indicated by arrows.

Since the fidelity of DSB repair can be a determinant of genome stability, we were interested in investigating the impact of progerin expression on the balance between PL and NHEJ and on other characteristics of DNA repair. To do so, we stably transfected cell line 13 with plasmid pEGFP-D50 lamin A which drives expression of GFP-progerin. Three derivatives of line 13 were identified that stably express GFP-progerin and we named these clonal cell lines alpha, beta and gamma. Each of these derivatives of cell line 13 displayed nuclear GFP fluorescence and displayed a band on a western blot expected for the GFP-progerin fusion protein when an anti-GFP primary antibody was used (Figure 2). The GFP fluorescence and western blot also revealed that expression of GFP-progerin was not equal in the three derivatives of line 13, with alpha displaying the highest level of progerin expression and beta displaying the lowest level expression. Comparison of DNA end-joining events recovered from parent cell line 13 with events recovered from lines alpha, beta, and gamma would reveal progerin’s effect on DSB repair. (GFP, which tags progerin in our experiments, does not impact DSB repair [55]).

**Figure. 2.**
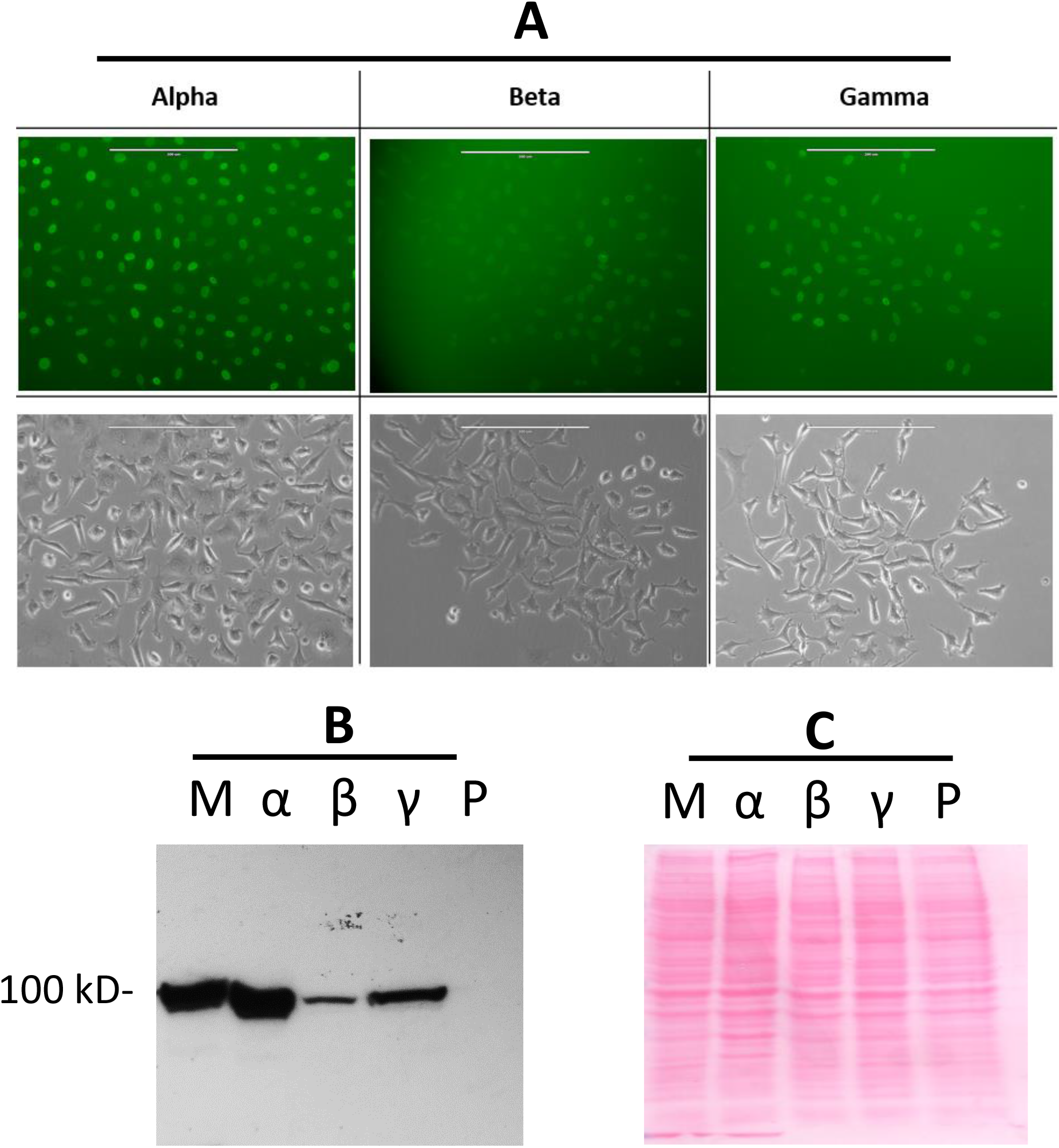
Expression of GFP-progerin in derivatives of cell line 13. **Panel A:** Nuclear fluorescence due to expression of GFP-progerin in cell lines alpha, beta, and gamma. For each cell line, a fluorescence microscopy image is presented above a light microscopy image of the same field of cells. **Panel B:** Western blot using a GFP-specific antibody to confirm expression of GFP-progerin in cell lines alpha, beta, and gamma (lanes α, β, γ respectively). Lane M displays a sample from cell line “pLB4-progerin” previously shown to express GFP-progerin at a level similar to progerin levels detected in cells derived from an HGPS patient [55] and this lane serves as a marker for GFP-progerin. Lane P displays a sample from parent cell line 13. Cell lines alpha, beta, and gamma each display the expected 100 kD band for GFP-progerin while the parent cell line 13 does not. Each lane contains 30 μg of total cellular protein. **Panel C:** Equal protein loading on the western blot is demonstrated by Ponceau staining of the membrane.

### Progerin expression shifts DNA end-joining away from PL and toward error-prone NHEJ

Cell line 13 and progerin-expressing derivative lines alpha, beta, and gamma were electroporated with pSce to induce DNA breakage and HAT^R^ colonies were recovered. As shown in Table 1, the colony frequencies for alpha, beta, and gamma were comparable to one another but were about two- to three-fold lower than the colony frequency for parent cell line 13.

**Table 1.** Recovery of DSB repair events

| **Table 1** |  |  |  |  |  |
| --- | --- | --- | --- | --- | --- |
| Recovery of DSB repair events |  |  |  |  |  |
| Cell line | Progerin expressed? | Number of experiments<sup>a</sup> | Total no. of cells plated (10<sup>5</sup>)<sup>b</sup> | Total no. of HAT<sup>R</sup> colonies<sup>c</sup> | HAT<sup>R</sup> colony frequency (10<sup>-3</sup>)<sup>d</sup> |
| Parent 13 | No | 3 | 6.0 | 2,199 | 3.7 |
| Alpha | Yes | 3 | 6.0 | 1,014 | 1.69 |
| Beta | Yes | 2 | 2.0 | 230 | 1.15 |
| Gamma | Yes | 2 | 4.0 | 668 | 1.67 |
| <sup>a</sup> Each experiment involved an electroporation of 5 x 10<sup>6</sup> cells with 20 µg of plasmid pSce to induce DSBs. |  |  |  |  |  |
| <sup>b</sup> Cells were plated at a density of 5 x 10<sup>4</sup> cells per 75 cm<sup>5</sup> flask after electroporation and subjected to HAT selection 48 hours after plating. Presented is the total number of cells plated among all electroporations for a given cell line. |  |  |  |  |  |
| <sup>c</sup> Total number of colonies arising under HAT selection. |  |  |  |  |  |
| <sup>d</sup> Total number of HAT<sup>R</sup> colonies divided by total number of cells plated. |  |  |  |  |  |

To determine the nature of recovered DSB repair events, genomic DNA was isolated from representative HAT^R^ clones recovered from cell lines 13, alpha, beta, and gamma. For each clone analyzed, a segment of DNA surrounding the DSB site was PCR-amplified, and the amplification products were sequenced. DNA sequence analysis allowed a distinction between PL and error-prone NHEJ. More specifically, as illustrated in Figure 1 and as described above, any clone that arose via a PL event will display a deletion of the 23 bp fragment of DNA between the two I-SceI sites and will retain the indicated 24 bp insert which includes a reconstituted I-SceI recognition site. In contrast, a clone that arose via error-prone NHEJ will display a distinct deletion or insertion restoring the tk gene reading frame.

Categorization of analyzed HAT^R^ clones as PL versus NHEJ is presented in Table 2. As indicated in Table 2, cell lines alpha, beta, and gamma each displayed a lower percentage of PL among recovered HAT^R^ clones compared to the results for parent cell line 13. Relative to parent cell line 13, the occurrence of PL was significantly reduced for alpha, beta, and gamma (p = 9.6 × 10^−5^, p = 1.2 × 10^−2^, and p = 5.8 × 10^−3^, respectively, by two-sided Fisher exact tests). The frequencies of occurrence of PL events among lines alpha, beta, and gamma were not statistically different from one another. These results suggested that expression of progerin reduces the likelihood of a DSB being repaired by PL. Interestingly, the magnitude of the deficit in PL correlated with the level of progerin expression, with cell line alpha displaying the greatest reduction in PL and cell line beta displaying the least reduction in PL.

**Table 2.** Categorization of DNA end-joining events

| **Table 2** |  |  |  |  |
| --- | --- | --- | --- | --- |
| Categorization of DNA end-joining events |  |  |  |  |
| Cell line | Progerin expressed? | Repair events analyzed<sup>a</sup> | PL<sup>b</sup> | NHEJ<sup>c</sup> |
| Parent 13 | No | 19 | 16 | 3 |
| Alpha | Yes | 31 | 8 | 23 |
| Beta | Yes | 25 | 11 | 14 |
| Gamma | Yes | 24 | 10 | 14 |
| <sup>a</sup> Total number of HAT<sup>R</sup> clones analyzed by DNA sequencing in the vicinity of the original site of DNA breakage. |  |  |  |  |
| <sup>b</sup> Total number of events retaining the 24 bp insert shown in Figure1 indicative of PL. The fraction of PL among end-joining events was significantly lower in cell lines alpha, beta, and gamma compared to 13 ($p = 9.6 \times 10^{-5}$, $p = 1.2 \times 10^{-2}$, and $p = 5.8 \times 10^{-3}$, respectively, by two-sided Fisher exact tests). | | | | |
| <sup>c</sup> Total number of events displaying sequence alterations distinct from PL. |  |  |  |  |

A more complete analysis of HAT^R^ clones is presented in Table 3. As revealed by sequence analysis, clones expressing progerin carried out a variety of NHEJ events, with deletion sizes ranging in size from 2 bp to as large as 47 bp. There were also five clones that arose due to the insertion of a single nucleotide (A) rather than deletion. All recovered HAT^R^ clones restored the correct tk reading frame. The deletion sizes associated with end-joining in progerin-expressing lines alpha, beta, and gamma appeared to be greater than deletion sizes recovered from parent 13. For example, if we consider the number of recovered NHEJ events with a deletion size exceeding 5 bp, there were significantly more such events among HAT^R^ colonies recovered from the progerin-expressing cells compared to parent cell line 13 (p = 0.036 by a two-sided Fisher exact test), and these larger deletions would be expected to be detrimental to cellular functions.

**Table 3.**
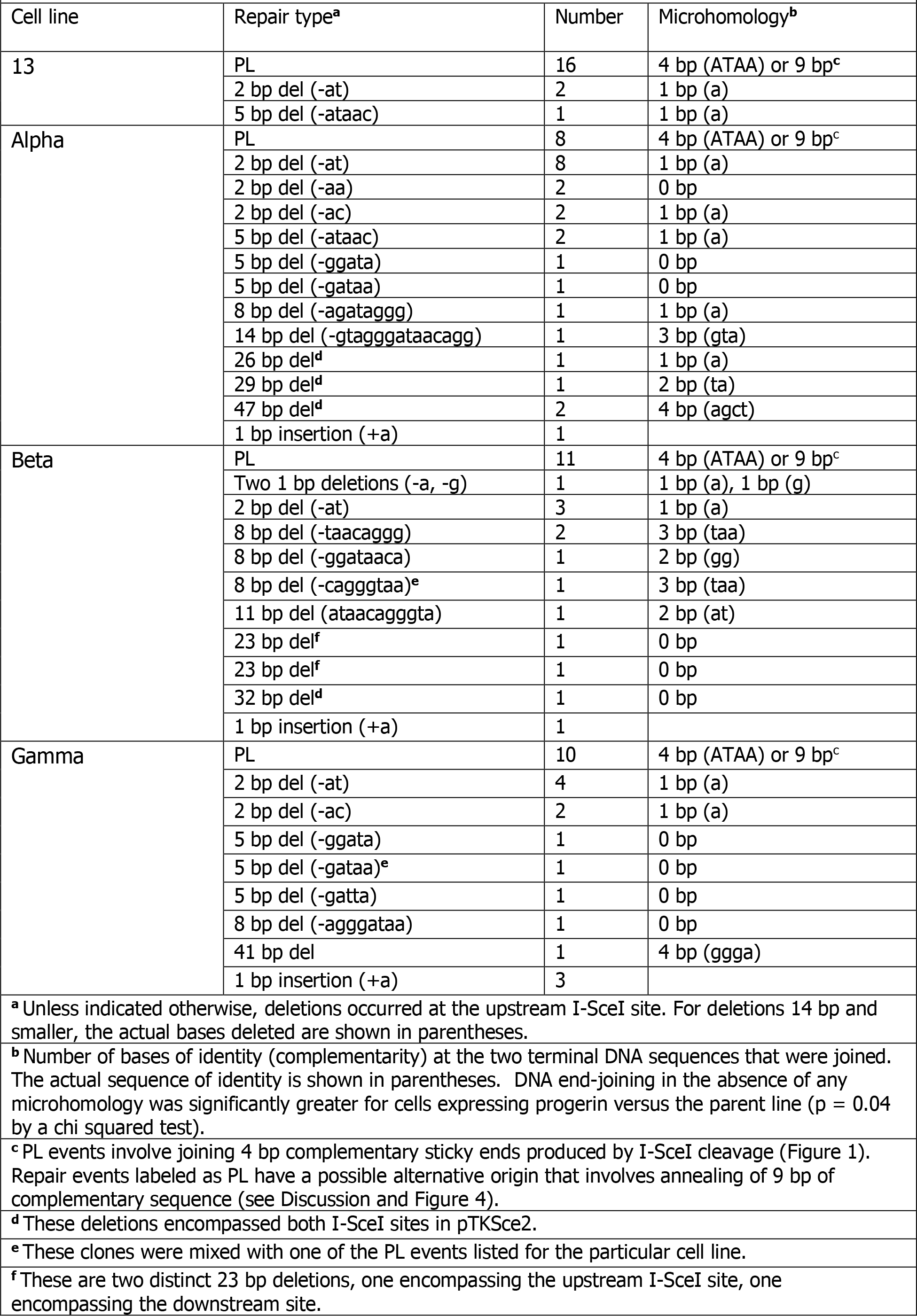
Analysis of DNA end-joining events

There was no notable requirement for substantial microhomology at the sites of DNA end-joining among progerin-expressing cells. Indeed, at least eleven clones from a total of 79 events recovered from alpha, beta and gamma arose from end-joining events involving no microhomology. In contrast, none of the events recovered from parent line 13 involved no microhomology. The difference in occurrence of DNA end-joining in the absence of microhomology was significantly different for cells expressing progerin versus the parent line (p = 0.023 by a chi squared test). Collectively, the reduction of PL events and the increase in end-joining events involving no microhomology in cells expressing progerin suggested a progerin-induced decrease in homology usage and/or recognition during DSB repair.

### Progerin expression does not enable homeologous recombination

Since the results presented above suggested that progerin expression resulted in a reduced usage of terminal sequence homology during DNA end-joining, we were curious to learn if progerin expression would lower the homology requirements of DSB repair via recombination. We therefore asked whether progerin expression would allow recombination between imperfectly matched sequences, also referred to as homeologous recombination (HeR). To address this issue, we made use of cell line HOME1-24 [60], which contains a single stably integrated copy of substrate pHOME (Figure 3).

**Figure 3.**
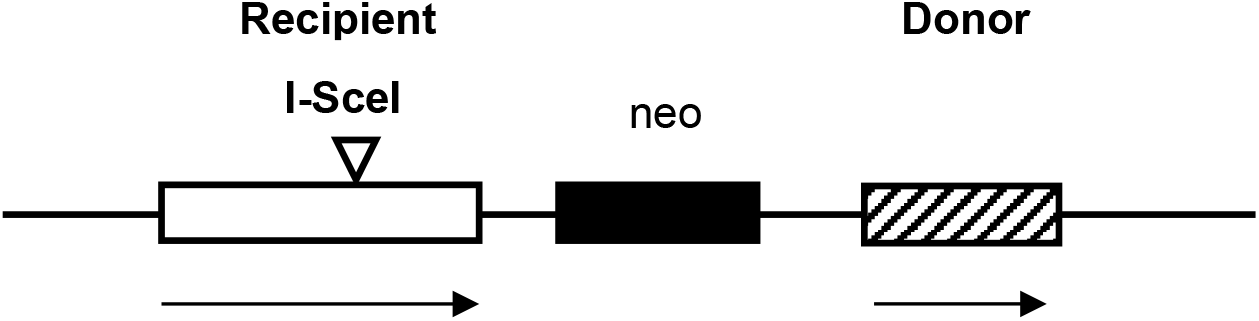
Substrate pHOME [60] for studying HeR. The “recipient” is a herpes simplex virus type one tk gene disrupted by an oligonucleotide containing an I-SceI recognition site. The “donor” is a fragment of the herpes simplex virus type two tk gene missing 30% of the tk coding region at the 5′ end. The donor and recipient display about 20% overall sequence divergence with scattered mismatches. Both crossovers and non-crossovers are potentially recoverable under HAT selection following HeR between the recipient and donor.

Substrate pHOME contains a “recipient” herpes simplex virus type one tk gene rendered nonfunctional by insertion of a 30-bp oligonucleotide containing the 18 bp recognition site for endonuclease I-SceI. Also included in pHOME is a closely linked “donor” sequence that shares about 80% nucleotide identity (20% divergence) overall with the recipient. The donor sequence on pHOME is a nonfunctional fragment of the herpes simplex type 2 tk gene missing 30% of the tk coding region at the 5′ end.

Substrate pHOME was designed as a reporter or HeR and NHEJ. Both crossovers and non-crossovers could potentially be recovered among any HeR events that might occur. Two derivatives of cell line HOME 1-24 that stably express GFP-progerin were isolated following transfection of HOME 1-24 with pEGFP-D50 lamin A. The progerin-expressing cell lines were named lines #1 and #5.

Lines #1 and #5 were electroporated with pSce and HAT^R^ colonies were selected (Table 4). Interestingly, the frequencies of recovery of HAT^R^ colonies for progerin-expressing cell lines #1 and #5 were about three-fold lower than the colony frequency for parent cell line HOME1-24, which was similar to the reduction in colony recovery observed for cell lines alpha, beta, and gamma relative to their parent cell line 13 (Table 1). Twelve HAT^R^ colonies from line #1 and 13 HAT^R^ colonies from line #5 were recovered and analyzed by PCR amplifying DNA sequence surrounding the DSB site followed by DNA sequencing (Table 4). One colony from cell line #5 was identified as an unequivocal homeologous recombinant based on the transfer of mismatched bases from the donor to recipient, with the balance arising from NHEJ. The sole HeR recombinant was a non-crossover event with a gene conversion tract including mismatched bases from the donor sequence mapping 30 bp upstream from the I-SceI cut site through 71 bp downstream from the cut site. The gene conversion tract encompassed 13 single base mismatches between donor and recipient. In previously published work with parent cell line pHOME 1-24 which does not express progerin, 2 out of 73 events DSB repair events recovered were homeologous gene conversion events encompassing 10 and 13 single base mismatches events, the balance of events being NHEJ [60]. Our results therefore provided no evidence that expression of progerin enabled HeR as a means of DSB repair.

**Table 4.** Recovery of homeologous recombination events

| **Table 4** |  |  |  |  |  |
| --- | --- | --- | --- | --- | --- |
| Recovery of homeologous recombination events |  |  |  |  |  |
| Cell line | Progerin expressed? | HAT<sup>R</sup> colony frequency (10<sup>-3</sup>) | HAT<sup>R</sup> colonies analyzed | NHEJ | HeR |
| #1<sup>a</sup> | Yes | 2.4 | 12<sup>b</sup> | 12 | 0 |
| #5<sup>a</sup> | Yes | 2.6 | 13<sup>b</sup> | 13 | 1 |
| HOME 1-24<sup>c</sup> | No | 7.8 | 73 | 71 | 2 |
| <sup>a</sup> Cell lines #1 and #5 are progerin-expressing derivatives of cell line HOME 1-24 [60] which contains a single integrated copy of pHOME (Figure 3) |  |  |  |  |  |
| <sup>b</sup> For each of cell lines #1 and #5, a single electroporation of 5 x 10<sup>6</sup> cells with pSce was performed and all recovered HAT<sup>R</sup> colonies were analyzed. |  |  |  |  |  |
| <sup>c</sup> Data for cell line HOME 1-24 is from [60] and represents collective results for four electroporations of 5 x 10<sup>6</sup> cells each. |  |  |  |  |  |

To assess whether progerin expression may enable spontaneous HeR events in the absence of artificial DSB induction, we carried out a fluctuation test for the two progerin-expressing lines #1 and #5. For each cell line, ten subclones were isolated, grown independently to 20 million cells per subclone, and then plated into HAT medium separately. A total of 200 million cells per cell line for a grand total of 400 million cells for both cell lines were plated into HAT selection. No HeR events were recovered, and indeed not a single HAT^R^ segregant was recovered. We previously reported a similar lack of recovery of any spontaneous HeR events from a cell line that contained pHOME and that did not express progerin [60].

## DISCUSSION

Progerin expression from a mutated gene for the nuclear lamina protein lamin A causes the rare premature aging syndrome HGPS. Low level production of progerin brought about by alternative splicing of the lamin A gene in healthy individuals has also been implicated in the biology of normal aging. Since both premature and normal aging are associated with, and possibly caused by, alterations in DNA repair, it is of significant interest to gain an understanding of how progerin expression may impact DNA repair. We had previously shown [55] that progerin expression shifts DSB repair away from the accurate HR pathway and toward the error-prone NHEJ pathway. In the current work we used a model cell culture system to ask whether progerin expression alters the very nature of DNA end-joining.

We divided DNA end-joining into the two broad categories that we refer to as PL and NHEJ. We use the term PL to describe high-fidelity end-joining with no alteration to the joined terminal sequences, and we use the term NHEJ to describe all other end-joining events in which terminal nucleotides are added or deleted at the site of end-joining. The data presented in Tables 1 and 2 show that in cell lines alpha, beta, and gamma which express progerin, DNA end-joining was shifted away from PL and towards NHEJ relative to events occurring in the parent cell line 13 which does not express progerin. Since PL preserves DNA sequence, the observed progerin-induced reduction in PL as a means for end-joining would likely serve to destabilize the genome and, thus, plausibly be a contributing factor to aging. Reduction in DNA end-joining fidelity is apt to have a synergistic destabilizing impact on the genome when acting in combination with a progerin-induced reduction in DSB repair via accurate HR.

The magnitude of the deficit in PL correlated with the level of progerin expression, with cell line alpha displaying the greatest reduction in PL and cell line beta displaying the least reduction in PL. This suggests the possibility that the impact of progerin on DNA repair may be progerin dosage dependent. This raises the intriguing possibility that levels of progerin expression in healthy individuals may correlate with a corruption of DNA transactions and perhaps with health-related issues associated with to the normal aging process, or with the progression of aging itself.

Although we have used repair events recovered using construct pTKSce2 as a proxy for PL as depicted in Figure 1, it has not escaped our notice that we cannot formally rule out an alternative explanation for the origin of clones that retain the 24 bp insert shown in Figure 1. Rather than being a product of clean ligation of the two outer-most DNA termini produced by two I-SceI breaks, it is possible that clones labeled as PL may be produced by breakage at the upstream I-SceI site followed by end-resection and single-strand annealing as illustrated in Figure 4. This event would involve pairing of two nine bp complementary sequences. Although this type of event is distinct from end-joining with no change to terminal sequences, it does indeed represent a repair event that involves exposure, recognition, and utilization of terminal sequence match, as would PL. As such, both PL and the single-strand annealing event depicted in Figure 4 can be viewed as types of homology-directed repair that share features broadly with the initial homology search of HR. A deficiency in searching for and pairing of complementary sequences in response to a DSB may also be the essence of why HR is reduced in the presence of progerin. It is worth noting that regardless of whether progerin blocks the clean joining of ligatable DNA ends, or blocks the recognition and annealing of complementary sequences, or blocks both capabilities, the impact of progerin would be detrimental since both of these processes have vital roles in preserving the genome in the presence of damage.

**Figure 4.**
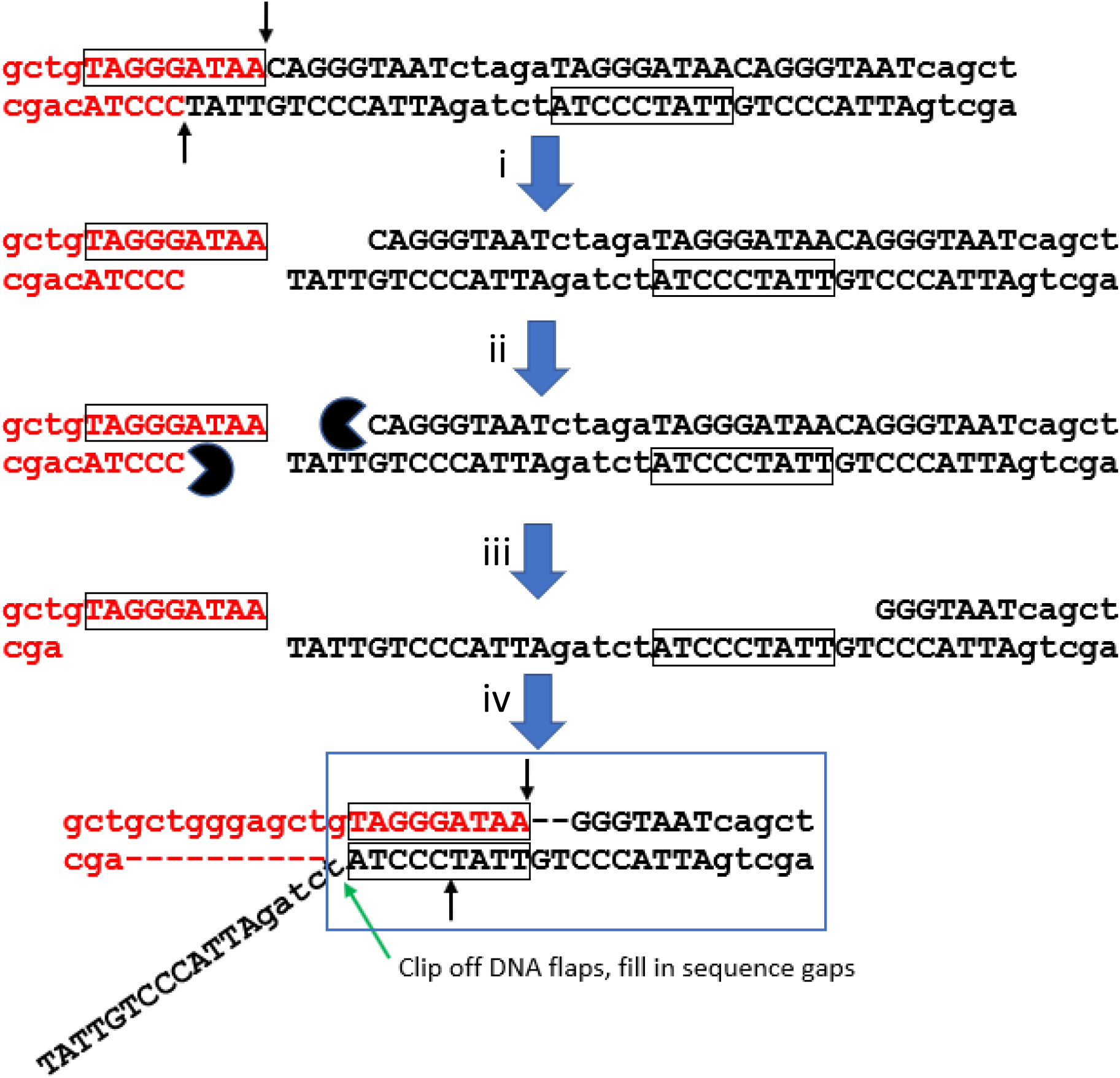
Alternative single-strand annealing origin of clones identified as products of PL. Clones indistinguishable from PL can potentially arise from the following process: i) the upstream I-SceI site in the 47 bp insert in pTKSce2 is cleaved by I-SceI; ii) 5’ ends of DNA strands are resected by exonuclease; iii) end-resection continues, exposing the two boxed 9 bp complementary sequences; iv) the boxed 9 bp sequences anneal. DNA flaps are clipped, and sequence gaps are filled to produce a clone that retains the 24 bp sequence (boxed in blue) that is also retained following PL. Short vertical arrows indicate sites of staggered cleavage within the I-SceI site.

The events identified as PL, recovered at high frequency from the original parent cell line 13, effectively result in loss of 23 bp at the site of genomic breakage and so it is possible to adopt the point of view that deletions associated with end-joining in parent line 13 actually tend to be larger than deletions in progerin-expressing lines alpha, beta, or gamma. In ongoing experiments, we have taken a PL clone recovered from cell line 13 and a PL clone recovered from cell line alpha and electroporated each with pSce to introduce a DSB at the I-SceI recognition site retained within the functional tk gene harbored in each PL clone. We then selected for cells resistant to trifluorothymidine (TFT^R^) [64] in order to recover clones that had lost tk function due to the induced DSB. (The loss-of-function assay should allow recovery of a less restrictive set of events than the original gain-of-function assay.) Following analysis of TFT^R^ clones, the genesis of some clones recovered remained unclear, but among the TFT^R^ clones that displayed a deletion at the I-SceI site the median deletion size in cells recovered from the cell line 13 PL clone was only 2 bp while the median deletion size in cells recovered from the alpha PL clone was 6 bp. The salient point is that cells recovered from the cell line 13 PL clone following the loss-of-function selection appeared to display deletion of fewer bp than the 23 bp loss seen in the predominant PL events recovered using the original gain-of-function selection. These observations suggest that parent cell line 13 likely has a propensity to delete very few terminal bp during end-joining at a DSB. This inference in turn indicates that, in the original experiment, the high recovery of events identified as PL in cell line 13 reflects a high rate of clean ligation of sticky DNA ends and/or a high rate of aligning and pairing of complementary sequences, but does *not* reflect an indiscriminately degradative pathway that simply deletes 23 bp of DNA duplex.

The data presented in Table 3 also reveal that DNA end-joining in the absence of any microhomology was a significantly more common occurrence in cells expressing progerin versus the parent line. This observation is consistent with the notion discussed above that annealing of terminal complementary bases is a less prominent feature of DSB repair in cells expressing progerin. The lesser involvement of terminal complementarity may be due to either an *impairment* in base-pairing between termini sequences or a reduced *dependency* of end-joining on terminal base-pairing in progerin-expressing cells. In a sense, these two different scenarios may figuratively be interpreted as progerin making DNA ends either “less sticky” or “more sticky,” respectively. The observations of larger NHEJ-associated deletions in progerin - expressing cells, the reduced use of HR in the presence of progerin [55], and the accumulation of DSBs in cells from HGPS patients seem most compatible with a progerin-associated suppression of interactions between terminal complementary bases of DNA, effectively making ends “less sticky” and more difficult to join. This impediment to end-joining could be a manifestation of the reported aberrant recruitment of nucleotide excision protein XPA to DSBs in cells expressing progerin [41]. It is believed that the mis-localization of XPA may functionally block normal DSB repair through stearic hindrance of the localization of DSB repair factors. The particular suppression of microhomology-mediated end-joining in cells expressing progerin might reflect a suppression of activity of polymerase theta which is known to play a key role in that process [65]. Various alterations to DSB repair which effectively make DNA ends less likely to be joined may ultimately be a manifestation of a progerin-induced change in chromatin structure. The overall reduced recovery of repair events that we recorded for progerin-expressing cell lines (Tables 1 and 4) may indeed reflect a general impediment to DNA end-joining.

Our observations that progerin prompted a decrease in PL and less involvement of microhomology in DNA end-joining evoked at least the possibility of an overall increased promiscuity of the DNA:DNA interactions that do occur in the presence of progerin and that might be brought about by impaired discrimination of sequence homology or complementarity. To explore this issue a little further, we were curious to learn whether progerin expression might reduce the homology requirements of HR and thus enable HeR. Our data with cell lines containing HeR construct pHOME and engineered to express progerin provided no evidence that progerin enabled HeR either following artificial DSB induction or spontaneously. These results are consistent with the view that progerin expression does not facilitate sloppy DNA:DNA interactions and does not lessen the dependency of DNA:DNA interactions on sequence homology or complementarity. Rather, in contrast, our collective results are consistent with a progerin-associated weakening of DNA:DNA interactions through an impediment to base pairing at DNA termini (i.e., DNA ends become “less sticky.”) We posit that this impediment leads to a reduced efficiency of DSB repair, reduced accuracy of end-joining, and fewer recombinational repair events, particularly crossovers, which require longer lived and more substantial interactions between DNA partners than does end-joining. It is possible that progerin exerts some of its impact on DNA repair by reducing the extent of DNA end-resection.

There are a variety of means by which a genome can become destabilized, and such destabilization may contribute to genetic disorders, including accelerated aging. Bloom syndrome, caused by a deficiency in Bloom helicase (BLM), is another rare genetic disorder that sometimes is classified as a premature aging syndrome. Bloom Syndrome, unlike HGPS, is associated with a high incidence of cancer. We had previously reported on the effects that deficiency in BLM has on DNA transactions [66, 67]. We observed that BLM deficiency results in increased use of HR versus NHEJ, with an increase in crossover types of HR events specifically [66]. We also reported that BLM deficiency enables HeR [67]. Our observed impacts of BLM deficiency are therefore basically the opposite of our observed impacts of progerin expression. To return to our earlier analogy, we may thus describe BLM deficiency as making DNA ends perhaps “too sticky,” in contrast to progerin expression which makes DNA ends perhaps “not sticky enough.” When DNA loses its stickiness, as in HGPS, much damage remains unrepaired, and DSBs and perhaps other lesions persist. This condition can lead to continuous futile damage signaling, loss of cellular functions, and cellular demise in the form of senescence or death. When DNA ends become “too sticky,” as in Bloom Syndrome, a cell may carry out reckless repair that may eliminate DNA breaks and allow a cell to continue to survive for some time, but at the expense of genetic rearrangements and mutation. These genetic alterations ultimately can lead to an accumulation of cellular dysfunctions, which in turn may lead to an accelerated aging phenotype and cancer. Thus, two seemingly antithetical alterations in DNA repair, that is, too little versus too much of a “good thing,” can lead to syndromes that effectively shorten life. Such observations illustrate the delicate balance that must be maintained in cellular pathways to preserve the health and well-being of the individual.

Life is quite complex, and DNA repair pathways are but one aspect of cell maintenance that plays a vital role in the biology of aging. To what degree changes or imbalances in DNA repair play causative roles in aging, are consequences of aging, or are a mixture of both, is at present very difficult to dissect. Additionally, we must remain cognizant that additional factors, including sex and nutritional status, undoubtedly influence the biology of aging, laminopathies in general, and indeed all biological processes. The interplay between such factors and health outcomes warrants much consideration and investigation. By illuminating the impacts that progerin expression or other genetic changes have on cellular and organismal well-being, and by probing the complex interplay between genetics and environment in the broadest sense, we will be in a better position to develop tools to improve the span and quality of life.

## Data Availability

Cell lines and plasmids are available upon request. The authors affirm that all data necessary for confirming the conclusions of the article are present within the article, figures, and tables.

## Acknowledgments

The authors are very thankful to Dr. Percy “Logan” Schuck and Dr. Jason Stewart for their excellent and generous assistance with western blotting. We also thank Dr. Katie Kathrein for so kindly allowing use of equipment that greatly facilitated our work, and we thank Zane Joudeh for excellent technical assistance.

## References

[1] M.S. Ahmed, S. Ikram, N. Bibi, A. Mir, Hutchinson-Gilford progeria syndrome: a premature aging disease, Mol. Neurobiol. 55 (2018) 4417–4427, https://doi.org/10.1007/s12035-017-0610-7. Epub 2017 Jun 28. PMID: 28660486

[2] E. Sonoda, H. Hochegger, A. Saberi, Y. Taniguchi, S. Takeda, Differential usage of non-homologous end-joining and homologous recombination in double-strand break repair, DNA Repair (Amst.) 5 (2006) 1021–1029.

[3] J. San Filippo, P. Sung, H. Klein, Mechanism of eukaryotic homologous recombination, Annu. Rev. Biochem. 77 (2008) 229–257.

[4] B. Pardo, B. Gómez-González, A. Aguilera, DNA double strand break repair: how to fix a broken relationship, Cell Mol. Life Sci. 66 (2009) 1039–1056.

[5] M. Shrivastav, L.P. De Haro, J.A. Nickoloff, Regulation of DNA double-strand break repair pathway choice, Cell Res. 18 (2009) 134–147.

[6] W.D. Heyer, K.T. Ehmsen, J. Liu, Regulation of homologous recombination in eukaryotes, Annu. Rev. Genet. 44 (2010) 113–139.

[7] E.M. Kass, M. Jasin, Collaboration and competition between DNA double-strand break repair pathways, FEBS Lett. 584 (2010) 3703–3708.

[8] J.R. Chapman, M.R. Taylor, S.J. Boulton, Playing the end game: DNA double-strand break repair pathway choice, Mol. Cell 47 (2012) 497–510, https://doi.org/10.1016/j.molcel.2012.07.029.

[9] A.A. Goodarzi, P.A. Jeggo, The repair and signaling responses to DNA double-strand breaks, Adv. Genet. 82 (2013) 1–45, https://doi.org/10.1016/B978-0-12-407676-1.00001-9.

[10] T. Iyama, D.M. Wilson III, DNA repair mechanisms in dividing and non-dividing cells, DNA Repair (Amst.) 12 (2013) 620–636, https://doi.org/10.1016/j.dnarep.2013.04.015.

[11] R. Scully, A. Panday, R. Elango, N.A. Willis, DNA double-strand break repair-pathway choice in somatic mammalian cells, Nat. Rev. Mol. Cell Biol. 20 (2019) 698–714, https://doi.org/10.1038/s41580-019-0152-0.

[12] G. Luo, I.M. Santoro, L.D. McDaniel, I. Nishijima, M. Mills, H. Youssoufian, H. Vogel, R.A. Schultz, A. Bradley, Cancer predisposition caused by elevated mitotic recombination in Bloom mice, Nat. Genet. 26 (2000) 424–429.

[13] J.R. LaRocque, J.M. Stark, J. Oh, E. Bojilova, K. Yusa, K. Horie, J. Takeda, M. Jasin, Interhomolog recombination and loss of heterozygosity in wild-type and Bloom syndrome helicase (BLM)-deficient mammalian cells, Proc. Natl. Acad. Sci. U S A. 108 (2011) 11971–11976.

[14] A. Ciccia, and S.J. Elledge, The DNA damage response: making it safe to play with knives, Mol. Cell. 40 (2010) 179–204, https://doi.org/10.1016/j.molcel.2010.09.019. PMID: 20965415

[15] A. Piazza, and W.D. Heyer, Homologous Recombination and the Formation of Complex Genomic Rearrangements. Trends Cell Biol. 29 (2019) 135–149, https://doi.org/10.1016/j.tcb.2018.10.006. E-pub 2018 Nov 26. PMID: 30497856

[16] A.S. Waldman, R.M. Liskay, Differential effects of base-pair mismatch on intrachromosomal versus extrachromosomal recombination in mouse cells, Proc. Natl. Acad. Sci. USA 84 (1987) 5340–5344.

[17] A.S. Waldman, R.M. Liskay R.M, Dependence of intrachromosomal recombination in mammalian cells on uninterrupted homology, Mol. Cell. Biol. 8 (1988) 5350–5357.

[18] T. Lukacsovich, A.S. Waldman, Suppression of intrachromosomal gene conversion in mammalian cells by small degrees of sequence divergence, Genetics 151 (1999) 1559–1568.

[19] Y. Lin, T. Lukacsovich, and A.S. Waldman, Multiple Pathways for Repair of DNA Double-strand Breaks in Mammalian Chromosomes, Mol. Cell. Biol. 19 (1999) 8353–8360.

[20] A.J. Bishop, R.H. Schiestl, Role of homologous recombination in carcinogenesis, Exp. Mol. Pathol. 74 (2003) 94–105.

[21] J.E. Eyfjord, S.K. Bodvarsdottir, Genomic instability and cancer: networks involved in response to DNA damage, Mutat. Res. 592 (2005) 18–28.

[22] R.D. Kennedy, A.D. D’Andrea, DNA repair pathways in clinical practice: lessons from pediatric cancer susceptibility syndromes, J Clin. Oncol. 24 (2006) 3799–3808.

[23] V. Gorbunova, A. Seluanov A. Making ends meet in old age: DSB repair and aging, Mech. Ageing Dev. 126 (2005) 621–628.

[24] K.J. Kyng, V.A. Bohr, Gene expression and DNA repair in progeroid syndromes and human aging, Ageing Res. Rev. 4 (2005) 570–602.

[25] D.B. Lombard, K.F. Chua, R. Mostoslavsky, S. Franco, M. Gostissa, F.W. Alt, DNA repair, genome stability, and aging, Cell 120 (2005) 497–512.

[26] V. Gorbunova, A. Seluanov, Z. Mao, C. Hine, Changes in DNA repair during aging, Nucleic Acids Res. 35 (2007) 7466–7474.

[27] A.A. Freitas, J.P. de Magalhães, A review and appraisal of the DNA damage theory of ageing, Mutat. Res. 728 (2011) 12–22.

[28] V. Tiwari, D.M. Wilson 3rd, DNA damage and associated DNA repair defects in disease and premature Aging, Am. J. Hum. Genet. 105 (2019) 237–257. doi: https://doi.org/10.1016/j.ajhg.2019.06.005.

[29] K. Ren, S.P. de Ortiz, Non-homologous DNA end joining in the mature rat brain, J Neurochem. 80 (2002) 949–59.

[30] J. Vijg, M.E.T. Dolle, Large genome rearrangements as a primary cause of aging, Mech. Ageing Dev. 123 (2002) 907–915.

[31] A. Seluanov, D. Mittelman, O.M. Pereira-Smith, J.H. Wilson, V. Gorbunova, DNA end joining becomes less efficient and more error-prone during cellular senescence. Proc. Natl. Acad. Sci. U S A. 101 (2004) 7624–7629.

[32] C.Z. Bachrati, I.D. Hickson, RecQ helicases: suppressors of tumorigenesis and premature aging, Biochem. J. 374 (2003) 577–606.

[33] M. O’Driscoll, P.A. Jeggo, The role of double-strand break repair - insights from human genetics, Nat. Rev. Genet. 7 (2006) 45–54.

[34] R.M. Brosh Jr., V.A. Bohr, Human premature aging, DNA repair and RecQ helicases, Nucleic Acids Res. 35 (2007) 7527–7544.

[35] C. López-Otín, M.A. Blasco, L. Partridge, M. Serrano, G. Kroemer, The hallmarks of aging, Cell 153 (2013) 1194–1217, doi: https://doi.org/10.1016/j.cell.2013.05.039.

[36] R. Burla, M. La Torre, C. Merigliano, F. Vernì, I. Saggio, Genomic instability and DNA replication defects in progeroid syndromes, Nucleus, 9 (2018) 368–379, doi: https://doi.org/10.1080/19491034.2018.1476793

[37] K. Cao, C.D. Blair, D.A. Faddah, J.E. Kieckhaefer, M. Olive, M.R. Erdos, E.G. Nabel, F.S. Collins, Progerin and telomere dysfunction collaborate to trigger cellular senescence in normal human fibroblasts, J. Clin. Invest. 121 (2011) 2833–2844.

[38] D. McClintock, D. Ratner, M. Lokuge, D.M. Owens, L.B. Gordon, F.S Collins, K. Djabali, The mutant form of lamin A that causes Hutchinson-Gilford Progeria is a biomarker of cellular aging in human skin, PLoS ONE 2 (2007) e1269. doi: https://doi.org/10.1371/journal.pone.0001269

[39] P. Scaffidi, T. Misteli, Lamin A-dependent nuclear defects in human aging, Science 312 (2006) 1059–1063.

[40] V.V. Ashapkin, L.I. Kutueva, S.Y. Kurchashova, I.I. Kireev, Are there common mechanisms between the Hutchinson-Gilford Progeria Syndrome and Natural Aging? Front. Genet. 10 (2019) 455. doi: https://doi.org/10.3389/fgene.2019.00455

[41] Y. Liu, Y. Wang, A.E. Rusinol, M.S. Sinensky, J. Liu, S.M. Shell, Y. Zou, Involvement of xeroderma pigmentosum group A (XPA) in progeria arising from defective maturation of prelamin A, FASEB J. 22 (2008) 603–611.

[42] P.R. Musich, Y. Zou, Genomic instability and DNA damage responses in progeria arising from defective maturation of prelamin A, Aging 1 (2009) 28–37.

[43] R.N. Serio, Unraveling the mysteries of aging through a Hutchinson–Gilford progeria syndrome model, Rejuvenation Res. 14 (2011) 133–141.

[44] P.R. Musich, Y. Zou, DNA-damage accumulation and replicative arrest in Hutchinson–Gilford progeria syndrome, Biochemical Soc. Trans. 39 (2011) 1764–1769.

[45] I. Gonzalez-Suarez, A.B. Redwood, D.A. Grotsky, M.A. Neumann, E.H. Cheng, C.L. Stewart, A. Dusso, S. Gonzalo, A new pathway that regulates 53BP1 stability implicates cathepsin L and vitamin D in DNA repair, EMBO J. 30 (2011) 3383–3396. doi: https://doi.org/10.1038/emboj.2011.225

[46] A.B. Redwood, S.M. Perkins, R.P. Vanderwaal, Z. Feng, K.J. Biehl, I. Gonzalez-Suarez, L. Morgado-Palacin, W. Shi, J. Sage, J.L. Roti-Roti, C.L., Stewart, J. Zhang, S. Gonzalo, A dual role for A-type lamins in DNA double-strand break repair, Cell Cycle 10 (2011) 2549–2560.

[47] H. Zhang, Z.M. Xiong, K. Cao, Mechanisms controlling the smooth muscle cell death in progeria via down-regulation of poly(ADP-ribose) polymerase 1, Proc. Natl. Acad. Sci. USA 111 (2014) E2261–E2270.

[48] S. Gonzalo, DNA damage and lamins, Adv. Exp. Med. Biol. 773 (2014) 377–399.

[49] I. Gibbs-Seymour, E. Markiewicz, S. Bekker-Jensen, N. Mailand, C.J. Hutchison, Lamin A/C-dependent interaction with 53BP1 promotes cellular responses to DNA damage, Aging Cell 14 (2015) 162–169. doi: https://doi.org/10.1111/acel.12258

[50] S. Gonzalo, R. Kreienkamp, DNA repair defects and genome instability in Hutchinson-Gilford Progeria Syndrome, Curr. Opin. Cell. Biol. 34 (2015) 75–83. doi: https://doi.org/10.1016/j.ceb.2015.05.007.

[51] X. Huang, Y. Pan, D. Cao, S. Fang, K. Huang, J. Chen, A. Chen, UVA-induced upregulation of progerin suppresses 53BP1-mediated NHEJ DSB repair in human keratinocytes via progerin-lamin A complex formation, Oncol. Rep. 37 (2017) 3617–3624, doi: https://doi.org/10.3892/or.2017.5603.

[52] C.D. Ragnauth, D.T. Warren, Y. Liu, R. McNair,T. Tajsic, N. Figg, R. Shroff, J. Skepper, C.M. Shanahan, Prelamin A acts to accelerate smooth muscle cell senescence and is a novel biomarker of human vascular aging, Circulation 121 (2010) 2200–2210.

[53] D. Constantinescu, A.B. Csoka, C.S. Navara, G.P. Schatten, Defective DSB repair correlates with abnormal nuclear morphology and is improved with FTI treatment in Hutchinson-Gilford progeria syndrome fibroblasts, Exp. Cell Res. 316 (2010) 2747–2759.

[54] B. Liu, J. Wang, K.M. Chan, W.M. Tjia, W. Deng, X. Guan, J.D. Huang, K.M. Li, P.Y. Chau, D.J. Chen, D. Pei, A.M. Pendas, J. Cadinanos, C. Lopez-Otin, H.F. Tse, C. Hutchison, J. Chen, Y. Cao, K.S. Cheah, K. Tryggvason, Z. Zhou, Genomic instability in laminopathy-based premature aging, Nat. Med. 11 (2005) 780–785.

[55] C.J. Komari, A. O. Guttman, S.R. Carr, T.L. Trachtenberg, E.A. Orloff, A.V. Haas, A.R. Patrick, S. Chowdhary, B.C. Waldman, A.S. Waldman, Alteration of genetic recombination and double-strand break repair in human cells by progerin expression, DNA repair 96 (2020) 102975. doi: https://doi.org/10.1016/j.dnarep.2020.102975

[56] A. Letsou, and R. M. Liskay, Effect of the molecular nature of mutation on the efficiency of intrachromosomal gene conversion in mouse cells. Genetics 117 (1987) 759–769.

[57] R.M. Liskay, J. L. Stachelek, and A. Letsou, Homologous recombination between repeated chromosomal sequences in mouse cells. Cold Spring Harbor Symp. Quant. Biol. 49 (1984) 183–189.

[58] P.S. Southern and P. Berg, Transformation of mammalian cells to antibiotic resistance with a bacterial gene under control of the SV40 early promoter. J. Mol. Appl. Genet. 1 (1982) 327–341.

[59] M.J. Wagner, J.A. Sharp, W.C. Summers, Nucleotide sequence of the thymidine kinase of herpes simplex virus type 1, Proc. Natl. Acad. Sci. U.S.A. 78 (1981) 1441–1445.

[60] V. Bhattacharjee, Y. Lin, B.C. Waldman, and A.S. Waldman, (2014) Induction of recombination between diverged sequences in a mammalian genome by a double-strand break, Cell. Mol. Life. Sci. 71 (2014) 2359–2371. doi: https://doi.org/10.1007/s00018-013-1520-0

[61] J.A. Smith, L.A. Bannister, V. Bhattacharjee, Y. Wang, B.C. Waldman, A.S. Waldman, Accurate homologous recombination is a prominent double-strand break repair pathway in mammalian chromosomes and is modulated by mismatch repair protein Msh2, Mol. Cell. Biol. 27 (2007) 7816–7827.

[62] W. Szybalski, E.H. Szybalska, and G. Ragni, Genetic studies with human cell lines. Natl. Cancer Inst. Monogr. 7 (1962) 75–88.

[63] K.M. Chapman, M.M. Wilkey, K.E. Potter, B.C. Waldman, A.S. Waldman, High homology is not required at the site of strand invasion during recombinational double-strand break repair in mammalian chromosomes, DNA Repair (Amst) 60 (2017) 1–8. doi: https://doi.org/10.1016/j.dnarep.2017.10.006.

[64] Y. Lin and A.S. Waldman, Capture of DNA sequences at double-strand breaks in mammalian chromosomes, Genetics 158 (2001) 1665–1674.

[65] K. Beagan and M. McVey, Linking DNA polymerase theta structure and function in health and disease, Cell. Mol. Life Sci. 73 (2016) 603–615. doi: https://doi.org/10.1007/s00018-015-2078-9.

[66] Y. Wang, K. Smith, B.C. Waldman, A.S. Waldman, Depletion of the Bloom syndrome helicase stimulates homology-dependent repair at double-strand breaks in human chromosomes, DNA Repair 10 (2011) 416–426.

[67] Y. Wang, S. Li, K. Smith, B.C. Waldman, A.S. Waldman, Intrachromosomal recombination between highly diverged DNA sequences is enabled in human cells deficient in Bloom helicase, DNA Repair 41 (2016) 73–84. doi: https://doi.org/10.1016/j.dnarep.2016.03.005.

